# A Unified Computational Framework for Deep Brain Stimulation at the Cellular and Network Levels

**DOI:** 10.64898/2026.07.02.736102

**Authors:** David Crompton, Luka Milosevic, Milad Lankarany

**Affiliations:** Institute of Biomedical Engineering, University of Toronto, Toronto, Ontario, Canada; Krembil Brain Institute, Toronto Western Hospital, Toronto, Ontario, Canada

**Author notes:** Membership list can be found in the Acknowledgments section.

## Abstract

Deep brain stimulation (DBS) has been demonstrated to be a successful therapeutic intervention for neurological disorders, yet the mechanisms underlying its effects on neuronal circuits remain incompletely understood. In this study, we propose a comprehensive phenomenological computational model that accounts for the impact of electrical stimulation parameters on neuronal circuits while incorporating experimentally-validated synaptic and cellular constraints. We investigate how DBS pulses modulate spiking activity in populations of homogeneous neurons representing stimulated nuclei, systematically examining the influence of circuitry architecture, including synaptic connectivity strength (weak vs. strong) and organization (sparse vs. rich). To characterize how DBS-modulated neuronal activity propagates through downstream networks, we develop a simple encoder that reveals distinct encoding patterns arising from different architectural configurations of stimulated nuclei. Furthermore, by connecting stimulated nuclei to recurrently connected neuronal populations, we examine the propagation of DBS-modulated neuronal synchrony across various circuit motifs. Our results demonstrate that three critical factors shape DBS-modulated neuronal activity: (a) the intrinsic synaptic and cellular properties of stimulated nuclei, (b) the architectural organization of stimulated nuclei in terms of synaptic strength and connectivity density, and (c) the circuit motifs formed by postsynaptic targets of stimulated nuclei. This unified model provides a mechanistic framework for understanding DBS representation and propagation in neuronal networks, offering insights that may inform optimization of stimulation parameters for clinical applications.

**Author summary:** Computational models of deep brain stimulation have proven to be supremely useful in disentangling the clinical benefits and adverse effects observed in the treatment of a variety of conditions. Despite this, the capacity for many of the existent computational models to account for micro/meso-circuit activation remains limited, as the major techniques rely on detailed characterization of tracts surrounding the DBS electrode, or depend on an non-physiologically constrained injected current intending to mimic the influence of electrical stimulation. The tract based methods only work for tracts that we have detailed characterization of, which are missing for many of the target structures, such as the basal ganglia. Given the restrictions of current methods we set out to define a phenomenological method that is applicable to as many simulation methods as possible, including those with missing details on tractography, while being able to readily integrate results of detailed simulations when available. Our approach is extensible and has examples implemented in some of the most popular computational neuroscience toolkits allowing for ready integration into existing network simulations. Further we demonstrate how this methodology supports interrogation of networks both for physiological responses but also computational dynamics, such as information multiplexing and delayed local evoked potentials.

## Introduction

Deep brain stimulation (DBS) is a neuromodulatory therapy used for a myriad of movement disorders [1–3], with a growing application to other psychiatric and neurological conditions [4–6]. Different hypotheses have been suggested to describe the method of action of DBS [7–9] and various computational models were developed to support those hypotheses [10–12]. Where, discounting non-neuronal effects, the primary effect is considered to be axonal activation. These axonal interactions include antidromic engagement, encapsulating a more global response to DBS. The localized impact of deep brain stimulation is also heterogeneous, dependent on the target nuclei [12], as well as the intranuclear location [13], which are thought to be driven by the specific organization of afferents as well as meso-network architecture. Deep brain stimulation also elicits different forms of short and long term synaptic plasticity [12, 14]. The complexities of the response to deep brain stimulation is what makes the versatility of computational models so useful as an exploratory and explanatory tool. The implementation of deep brain stimulation within computational models varies widely, we categorized DBS computational models in the literature based on two levels of descriptions, (1) detailed biophysical models of (2) abstract models. Detailed biophysical models of DBS were developed to study how physical characteristics of electrodes, stimulation parameters, and tissue substrates contribute to neural activities [10, 15, 16]. These methods allow for high accuracy characterization of neuronal engagement, accounting for non-faradaic/non-linear interactions, and the pulse waveform, which have been shown to substantially influence the resulting neural recruitment [11, 17, 18].

However, these methods depend on the accuracy and existence of a 3D representation of the neuronal structure [19], and these models have an immense computational burden, which makes it prohibitive to study how DBS modulates dynamics of a large population of neurons. Abstract models, consider the impact of DBS by some simplification, such as the injection of supra-threshold current [20, 21], or of some modified rate function [22]. The simplicity of this method, and lack of physical dependence, allows abstract methods to more thoroughly explore the response of large networks, and the range of responses in a more computationally tractable manner. Although abstract models can describe some aspects of DBS-induced changes (even at the firing rate level [22]), these abstractions can miss important effects such as antidromic activation [22], somatic-axonal decoupling [20, 21], or the spatial engagement shaped by stimulation settings. In this work we intend do develop a technique that can account for the varying impacts of deep brain stimulation, without loss of generalizability to more detailed or abstract systems.

Building on our previous computational models of DBS that capture various temporal dynamics of instantaneous firing rates of stimulated neurons in different sub-cortical regions (DBS targets) [12], we propose a unifying algorithmic framework that enables modelling the impact of DBS in any spiking neural network, conceptually described in [23] and implemented initially in [13]. Specifically, inspired by a previous work [24], we model the impact of DBS using direct synapse activation, or when missing a module, referred to as parrot neuron [25], to generate DBS-induced spikes on both afferents and efferents of stimulated neurons in a neuronal network, which allows for the heterogeneity and specificity of the response to stimulation [23]. The activation of afferents and efferents serves as an extension prior works which characterized the efferent activation [24], afferent activations [12], and antidromic activation [21]. The interaction of DBS with synaptic plasticity is incorporated as implemented in previous works [12, 21]. Which led to the model utilized in [13], integrating afferent, efferent, antidromic activation, and synaptic plasticity, which we set out to generalize in this work. We show that the use of parrot neuron preserves interactions of spike times of neurons and those generated by DBS, thereby enables accurate adjustments of synaptic dynamics altered by DBS. This simple abstraction to synaptic activations allows for a thorough investigation into the effects of deep brain stimulation on afferent, efferent, and antidromic connections. Additionally, compared to the current injection methods [20], and those that create artificial synapse pools activated by DBS [21] the plastic changes induced by DBS persist to future somatic/physiological activity.

Given the heterogeneity of response, connectivity, and dynamics in response to DBS the simulation study in this work is intended to demonstrate how plasticity and connectivity can shape information propagation. To showcase the generality of our DBS method, inspired by [26], our simulation study explores how deep brain stimulation both propagates through, and modulates a given circuit motif with a short term synapse dynamic. Wherein the different circuit motifs and plasticity dynamics represent different hypothetical circuits that may be simultaneously but differentially engaged by DBS. Where, as an formalization of our prior work [13], we showcase the effect on firing on both the afferent and efferent nuclei, as well as the local field potential. In our simulation study, we demonstrate that the impact of DBS on the firing rate and local field potential within different circuit motifs can be readily modelled and interrogated with our method. We anticipate that this work provides a standard modelling approach for studying how DBS alters dynamics of large neuronal populations. A representative figure of the features that this abstraction accounts for is shown in Figure 4.

## Materials and methods

In this work we describe a standard method of implementing Deep Brain Stimulation for spiking neural networks, as first implemented in [13]. The primary mode of action of this method is direct synaptic activation, applied to efferent [24], afferent [12], and antidromic connections [27]. Simulators like NEURON [28] allow for the delivery of event times directly to synapses, whereas other simulators such as NEST [25] and BRIAN [29] require the introduction of an intermediary process, referred to in NEST as, a parrot neuron. Parrot neurons serve to emit every spike they receive, their necessity derives from how some simulators do not allow for manual event delivery to synapses, thus intercepting the spike delivery with a ‘neuron’ that emits what it receives, allows us to emit both somatic spikes and those induced by DBS. Parrot neurons are to be placed preceding every synapse, refer to Figure 2 for visual representation.

**Fig 1.**
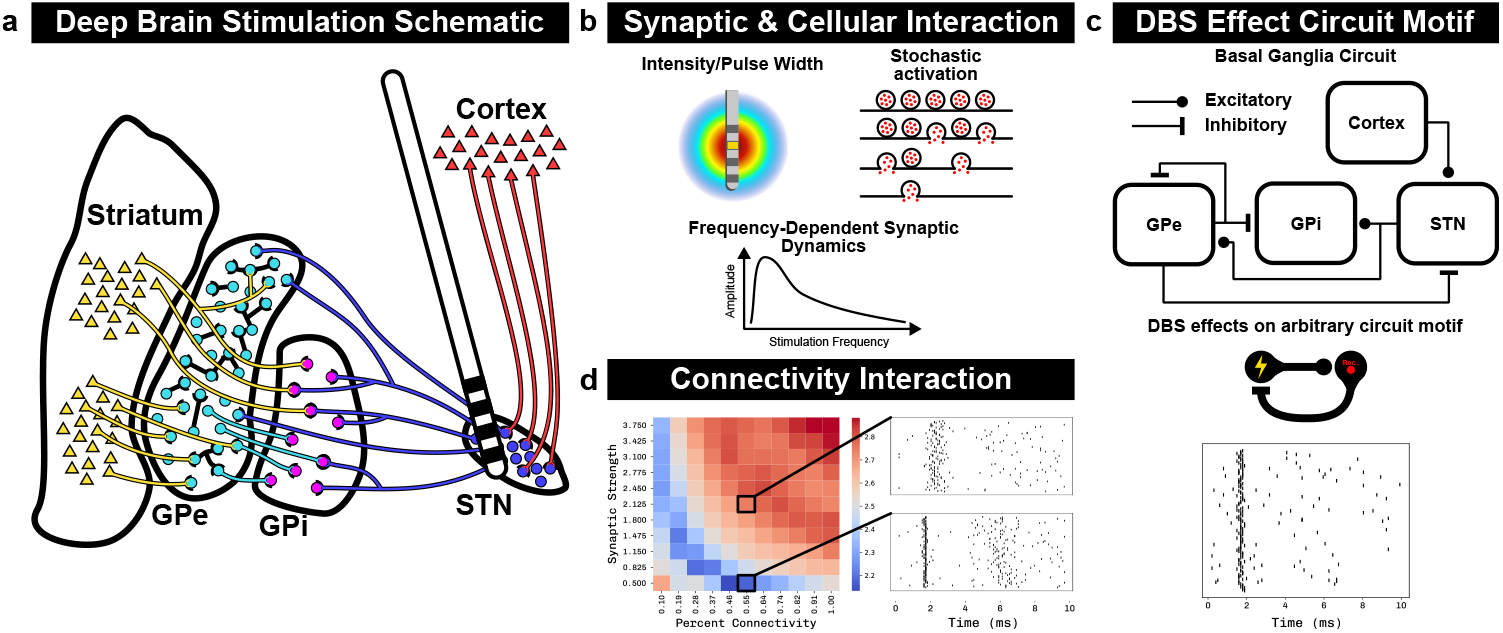
Deep Brain Stimulation Simulation framework. The motivation and underlying features of the abstraction of DBS. **a)** Schematic representation of a common network, the basal ganglia, where DBS is delivered. **b)** The cellular and synaptic features that an abstraction of DBS must account for, intensity recruitment, synaptic failure/stochasticity, and frequency dependent activity e.g. plasticity. **c)** The interaction of DBS with circuit architecture, feed-forward, lateral inhibition, recurrent inhibition. **d)** The meso-circuit properties, the interaction of DBS with connection sparsity and strength.

**Fig 2.**
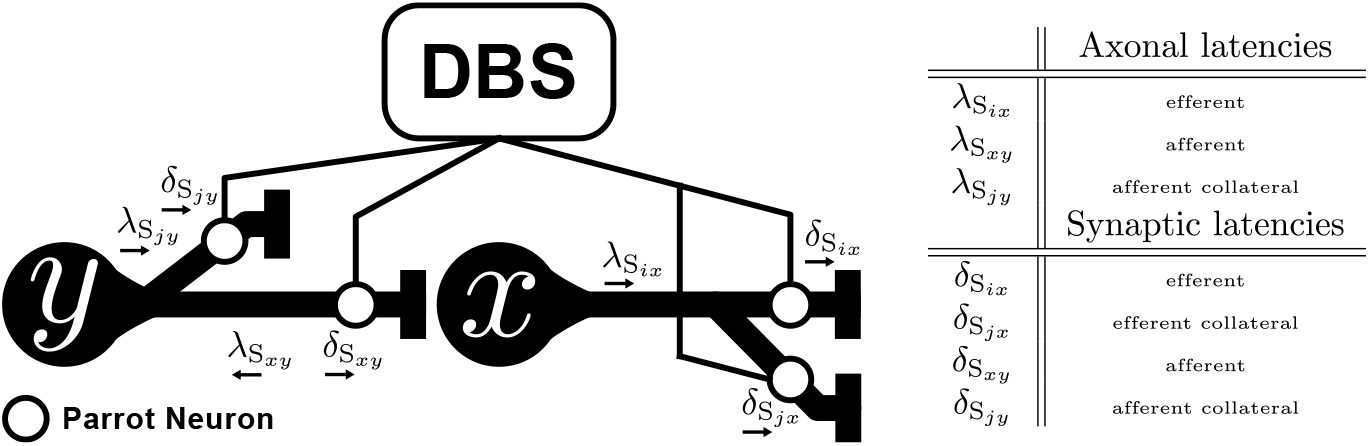
Visualization of placement of parrot neurons preceding all efferent and afferent synapses, and connectivity to the DBS spike generator. Axonal (*λ*) and synaptic (*δ*) latencies are depicted. Arrows below depict the propagation direction induced by DBS, where 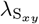 represents the antidromic propagation time

Once manual event delivery implemented, the following algorithm can be followed to implement DBS. The target population is maximal group that might be activated by stimulation, i.e. the nucleus that is stimulated, or the neurons within an area around the electrode. The neurons that are activated are determined by the function *Choose*, which may depends on the stimulation parameters, this function is application specific, variants are elaborated in the section. The probabilities [*p*_act_, *p*_eff_, *p*_aff_, *p*_anti_, 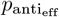] represent the chance for each DBS pulse to activate a given neuron or synapse. The only properties needed for each synapse, to determine the activation latency by DBS, are the axonal 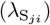 and synaptic 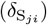, representing the delay needed for an action potential to propagate from the soma to the synapse, and the time for a synapse induce a post-synaptic potential respectively. How the latencies combine are depicted in Figure 2.

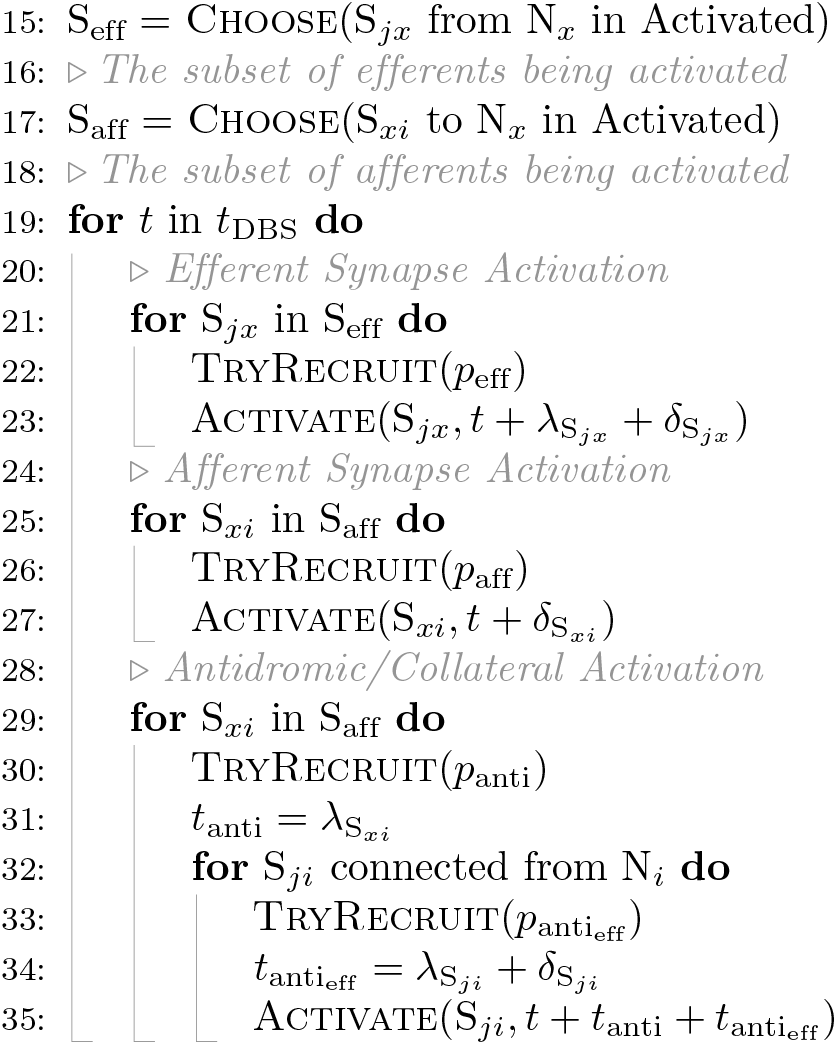
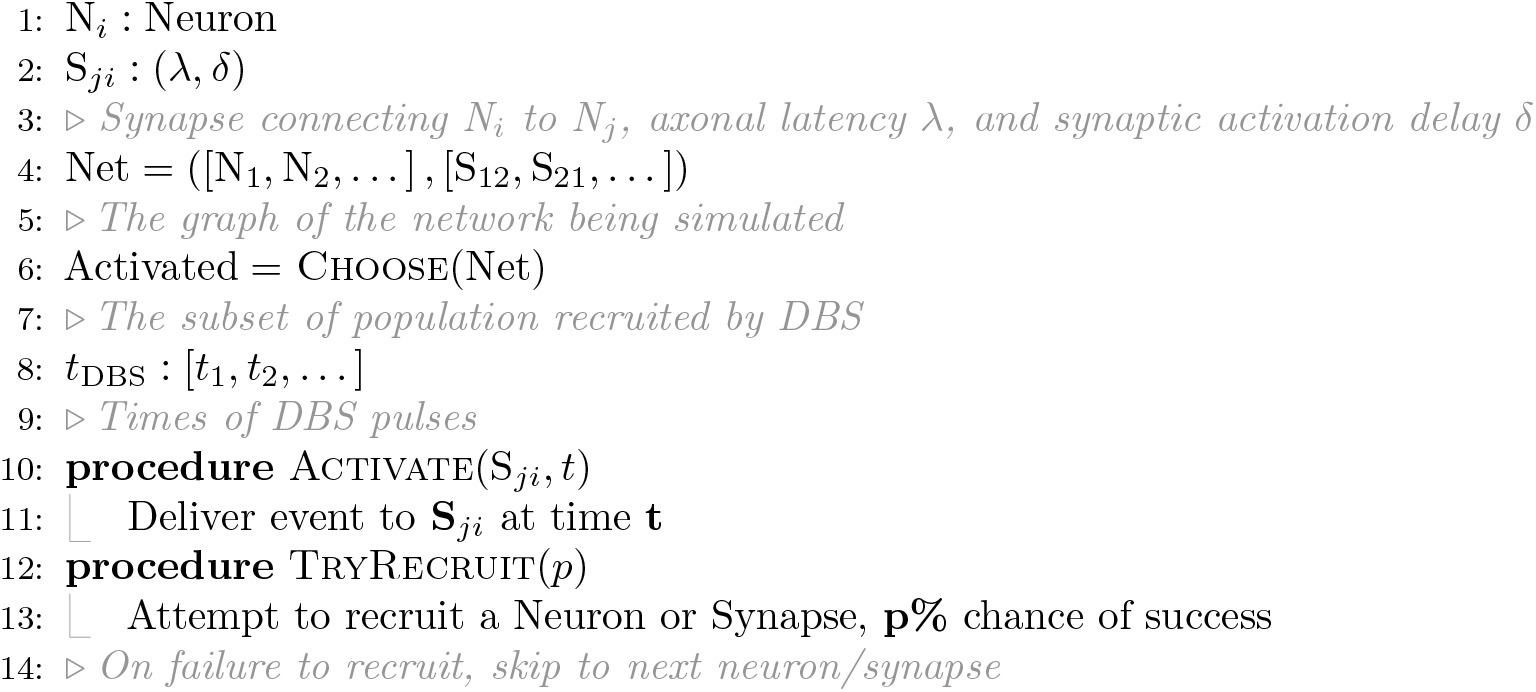

### Choice of activation

The *Choose* function represents an application specific function that determines the subset of neurons or synapses to be activated. For example, the subset of activated neurons (*Activated*) can be considered as the group of neurons encompassed by the Volume of Activated Tissue (VAT).

### Stimulation Parameter Effects

Parameters such as stimulation intensity, pulse-width, pulse shape, are intended to be considered by choice of *Choose* function. The sections below detail the considerations that should be made.

### Stimulation Intensity and Pulse-width

In consideration of the effect of intensity and pulse-width the *Choose* function may be constructed in a few ways, depending on the degree of abstraction desired. In the most abstract form of activation changes in stimulation intensity and pulse-width may correspond to a proportional increase in percent activation 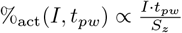, where *S*_*z*_ is the radius of the target population. Effects of pulse width, as well as pulse shape, e.g. symmetric/asymmetric, cathode or anode first can also be used to steer activation profiles (Figure 3). Anatomical features, such as fiber orientation can also influence the degree of recruitment for a given stimulation [9]. Antidromic propagation is described starting on Line 28 of the pseudocode. Antidromic activation especially relies on the *TryRecruit* behaviour to incorporate the action potential propagation failure [27]. Building on previous methods of estimating VAT derived from electric field estimates, either using point source (citation) or finite element modeling (citation) may be used, wherein the second derivative (citation) of the electric field may be used to determine what fraction of the population will be activated.

**Fig 3.**
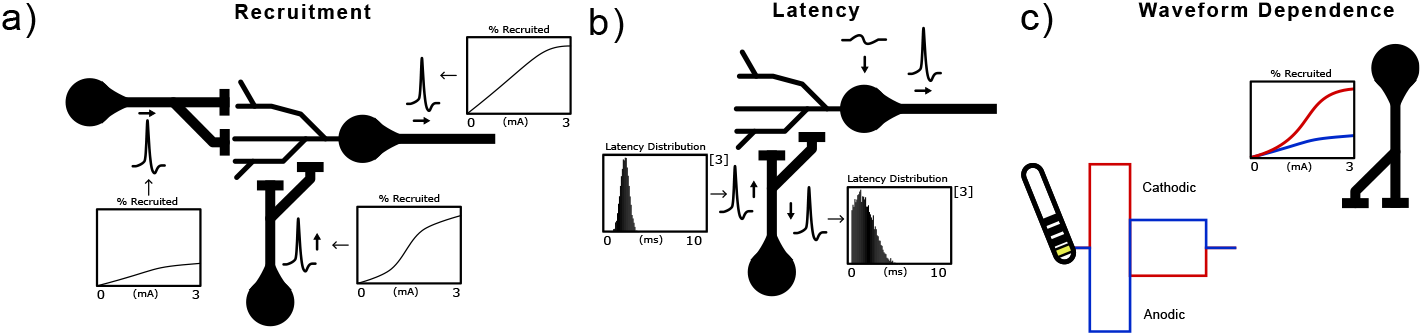
Stimulation recruitment considerations. **a)** Differential recruitment of fibers with respect to fiber properties and orientation, and DBS parameters. **b)** Latency modulation by stimulation parameters, fiber features, and orientation. **c)** Fiber recruitment influenced by stimulation pulse shape, anode/cathode first.

### Stimulation Frequency

Since this method relies on direct activation of the synapses in the model, it has a direct interplay with synaptic plasticity. Wherein, if a synapse has some underlying facilitation or depression, DBS will be shaped by the plasticity of the underlying synapse. To demonstrate this, we incorporate the Tsodyks Markram model of synaptic plasticity to showcase how short-term synaptic depression/facilitation effects impact of DBS (eqs. (4) to (7)).

### Synaptic Reliability

Since this method interacts on the synapse level, it allows for the additional characterization of the reliability of recruitment of each synapse. This is represented by the notion of a percent activation under the subset of the recruited synapses, where even if a given synapse is enveloped in the “volume of activated tissue” of the stimulation, it will not necessarily be recruited by stimulation, and can be tuned to what rate of recruitment is per pulse. This allows for more fine-grained interrogation of DBS on synaptic dynamics, as plasticity/vesicle dynamics change substantially with stochastic release [30].

### Model Equations

#### Neuron Model

The model neuron used in this work is the leaky integrate and fire (LIF) neuron, though the choice of neuron is not specific to the DBS abstraction.

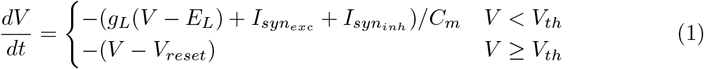

Where *V* is the voltage of the neuron model, *I*_*syn*_ is the excitatory and inhibitory inputs of the specific neuron, with *g*_*L*_ and *E*_*L*_ being the conductance and equilibrium potential of the leak current, and *C*_*m*_ being the membrane capacitance of the neuron. The voltage resets to *V*_*reset*_ when *V > V*_*th*_. Neuron parameters were *C*_*m*_ = 250 *pF, V*_*th*_ = *−*55 *mV, V*_*reset*_ = *−*60 *mV, g*_*L*_ = 16.6667 *nS,E*_*L*_ = *−*70 *mV, E*_*ex*_ = 0 *mV, E*_*in*_ = *−*80 *mV*,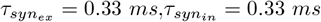

#### Synaptic Dynamics

Synaptic currents (*I*_*s*_*yn*) are given by the alpha kernel (Equation (3)), synaptic plasticity is represented by the phenomenological model of synaptic plasticity, the Tsodyks Markram model [31] in (eqs. (4) to (7)). Depression was implemented with *U* = 1,*u* = 1,*τ*_psc_ = 3 *ms,τ*_fac_ = 0 *ms,τ*_rec_ = 50 *ms*.

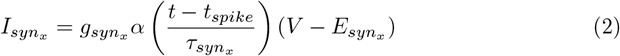

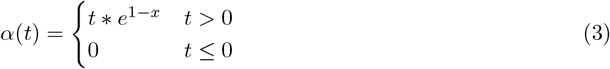

Where the maximal conductance of the synapse *g*_*synx*_ = *y* in the equations describing the short term-plasticity (Equations (4) to (7)). The rise and fall time of the post-synaptic kernel are controlled by 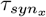, and the drive, excitatory or inhibitory is driven by the equilibrium potential 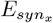.

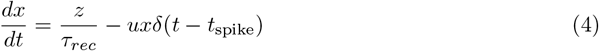

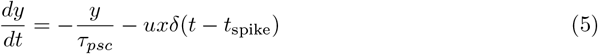

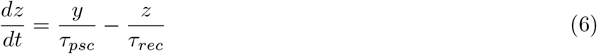

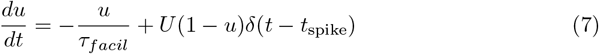

Where *u* is the initial quantity of neurotransmitter available changed by U on activation, and *u* is the change induced in x upon transmission. *τ*_*psc*_ is the width of the post-synaptic kernel, while *τ*_*fac*_, and *τ*_*rec*_ are the facilitation and recovery terms driving the short term dynamics.

### Simulation framework

The simulations presented herein are simulated in NEST [25], where neuronal populations are simulated with LIF neurons with alpha synapses, unless otherwise specified, equations eqs. (1) to (3). In addition to the otherwise specified connectivity, each population receives inputs from 10 excitatory, and 2 inhibitory from 10 *Hz* poisson spike generators. Unless otherwise specified, all synapses have an activation probability of 80%, and 80% of neurons were recruited.

### Results

### Deep brain stimulation Implementation

To demonstrate the effects of this DBS model, we constructed a simple feed forward recurrent network of two excitatory populations Figure 4. The efferent synapses are embedded with high frequency facilitatory dynamics (*U* = 0.001, *u* = 0, *τ*_fac_ = 50 *ms, τ*_rec_ = 5 *ms*), while the afferent synapses with high frequency depressive dynamics (*U* = 1,*u* = 1,*τ*_psc_ = 3 *ms,τ*_fac_ = 0 *ms,τ*_rec_ = 50 *ms*). We then simulated this network under DBS at 25 *Hz*, and 100 *Hz*, at both low and high intensity. The low and high intensity represented the percentage of neurons recruited by each DBS pulse, 20% and 80% respectively. The resultant raster plots of activity during stimulation of both populations is show in Figure 4, for each raster the bottom half represents the stimulated population, and the top half the efferent population.

**Fig 4.**
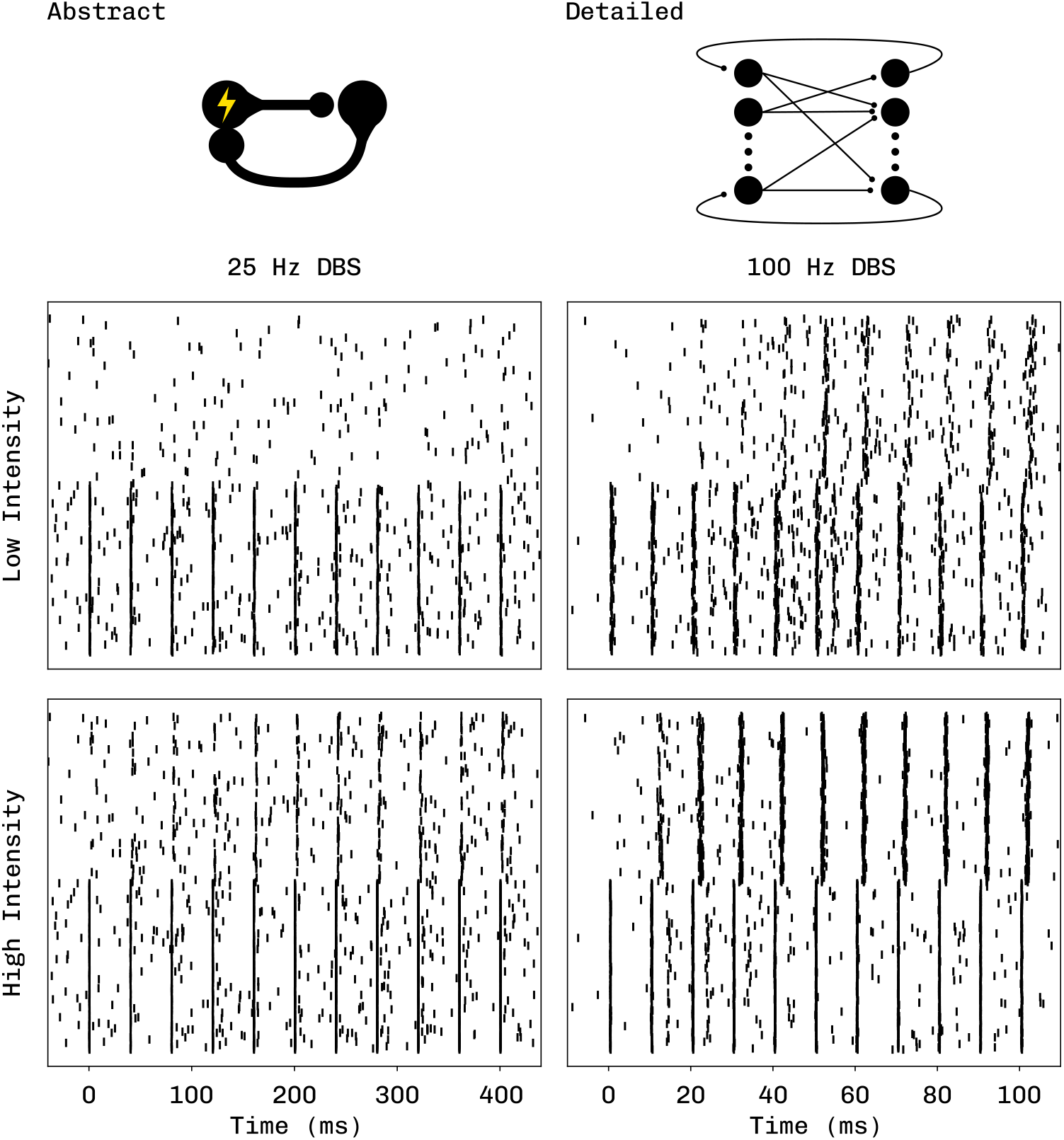
Deep Brain Stimulation Exemplary Motif. Spike rasters of the stimulated (bottom half) and efferent population (top half) are shown under three conditions for the first 10 pulses of stimulation. DBS parameters such as frequency and recruitment fidelity are shown in the columns and rows respectively.

The low frequency stimulation succeeds in recruiting afferents to induce activity in the stimulated population, while failing to sufficiently excite the efferent population at a low intensity. High frequency stimulation at a low intensity is able to, after a few pulses, entrain the efferent population due to the short term facilitation embedded in the model. At a high intensity there is an initial recurrent entrainment that persists for *∼* 5 pulses due to the embedded short term depression. The stimulation method fo DBS described functions independently of network size, as shown in Supplementary figure, the behaviour of stimulation extends when the number of neurons per layer is increased from 150 to 500 (Supplementary).

### DBS Representation in Feed Forward Circuits

Since DBS is often considered to disentangle axonal and somatic activity, we examine the propagation of activation induced by DBS while varying sparsity and connectivity strength, inspired by [32].

While the propagation synchrony varies similarly previously shown [32], we explore how information is propagated despite the precision in timing. We examine the frequency coding, as well as the baseline rate normalized frequency code with the same variation in sparsity and strength. While the rate increases proportionally to the increase in functional connectivity, either an increase in strength or number of connections, the normalized rate reaches a point of inflection after which DBS suppresses activity.

### Circuit Motif Influence on DBS

As a continuation of Figure 5 this method can be readily embedded within different circuit motifs. The DBS induced synchrony within different motifs is shown in Figure 7. The motifs used are a feed-forward network, feed-forward lateral inhibition, and feed-forward inhibitory recurrent connectivity. All synapses in each motif have no synaptic plasticity.

**Fig 5.**
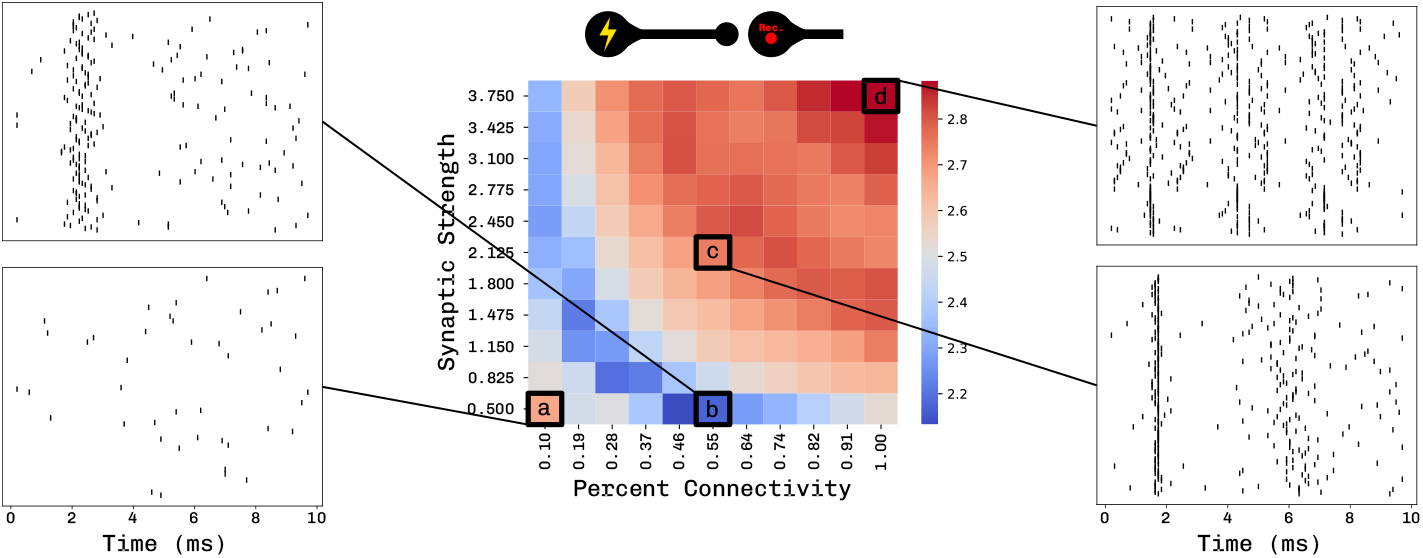
Stimulation-connectivity interaction. Influence of network sparsity and connectivity strength on the spiking activity in a feed-forward network model. The left figure represents the synchrony of spiking activity with respect to connection probability on the x-axis, and synapse strength on the y-axis. The rasters depict exemplary inter-stimulus interval raster plots of the first pulse of DBS on the efferent population, with lines indicating to the figure on the left what the sparsity and strength values were. **a** Low connectivity and low strength, **b** low strength and medium connectivity, **c** medium strength and medium connectivity, **d** high strength and high connectivity.

Temporal precision of DBS propagation is steered directly by the circuit motif surrounding the stimulated nuclei. Wherein the response for a specific set of connectivity parameters is different drastically different; the landscape of temporal precision as it relates to connectivity parameters is changed entirely, local and global optima are shifted.

### Local Field Potential Interaction

Extending the recurrent motif to have facilitation dominant synapses we examine low and high frequency stimulation, as well as low intensity high frequency stimulation. Here we observe that despite very similar raster profiles induced by stimulation, in this motif only high intensity high frequency stimulation can induce an apparent oscillation in the local field potential proxy (Figure 8, Figure 9). The evolution of the delayed evoked potential, the second peak, is shown in Figure 8.

The recurrent network with synaptic facilitation toward the efferent nuclei si able to induce an entrained oscillatory response that grows with the stimulation train. To compare and represent frequency and intensity dependent dynamics we simulate the average ISI response as observed in the same network stimulated at 10, 50, and 100 *Hz* and a low or high intensity. With the STP dynamics embedded a strong oscillatory-like response could only be observed at a high frequency and high intensity (Figure 9), lowering either the intensity or frequency substantially reduced the delayed peak at *∼* 8 *ms*.

## Discussion

In this work we described a comprehensive method by which one can implement the impact of DBS in any spiking neural network. By directly on afferent and efferent synapse populations we are able to account for a variety of features thought to contribute to the impact of stimulation, such as recruitment probability, and latency of activation (Figure 2). To showcase how the effects of short term plasticity interact with DBS we constructed both a feed-forward and recurrent network, and stimulated at different frequencies and intensities to probe the response, both in the spiking activity and proxy for local field potential. Within this we demonstrate how low intensity DBS may fail to recruit the efferent populations at low frequencies when under the influence of short-term facilitation at the higher frequencies. Similarly, how at high intensity and high frequency the recurrent recruitment in the stimulated population may be lost from short term depression. Prior methods of simulating the interaction of DBS with short term plasticity did so by introducing a time-varying current that would be injected to simulate the effect of DBS [21, 22]. By contrast, our method does not necessitate the creation of currents specific to DBS, as it acts solely on existent synapses, thereby inheriting their synaptic dynamics.

Since deep brain stimulation is often thought to act as an information lesion, we investigated how DBS impacts different motifs on the ability encode information. As illustrated in Figure 3-5 the capacity for the circuit to support the propagation of DBS, and in turn how readily it is able to lesion information processing is highly sensitive to both parameters and motif. In the simplest feed forward case (Figure 4) we see that how DBS effects efferent nuclei, and when DBS optimally interferes, or replaces the encoded information depends greatly on the modality of encoding. We depict three different encoding schemes, temporal precision, rate coding, and normalized rate coding. For temporal precision and normalized rate encoding there is a non-monotonic relationship to encoding with respect to connectivity strength and probability. Intuitively, the pure rate encoding scheme follows a monotonic relationship, as in a feed forward network the increase in connectivity directly corresponds to an increased drive by DBS. When comparing the temporal precision across motifs however, we diverge from [26], as inhibition stabilizes the spike propagation, especially in the lateral inhibition case [33]. The intent of these simulations is to showcase that the effect of DBS is shaped strongly by the properties of the network being stimulated, for which this method provides easy interrogation. An exemplary interrogation is of a recurrent network representing part of the basal ganglia, as it pertains to features such as Evoked Recurrent Neural Activity (ERNA) [13] and Delayed Local Evoked Potentials [15] known to be related to clinical improvement in conditions such as Parkinson’s disease and Dystonia [13, 34]. These responses are observed as an oscillatory response to stimulation in the local field potential, which grows with frequency and intensity. For Figures 6-7 the recurrent circuit has short term facilitation between the nuclei, and we can directly see the interplay of the plasticity dynamics, wherein the apparent oscillation is strengthened by the facilitation at higher frequencies and the intensity of stimulation.

**Fig 6.**
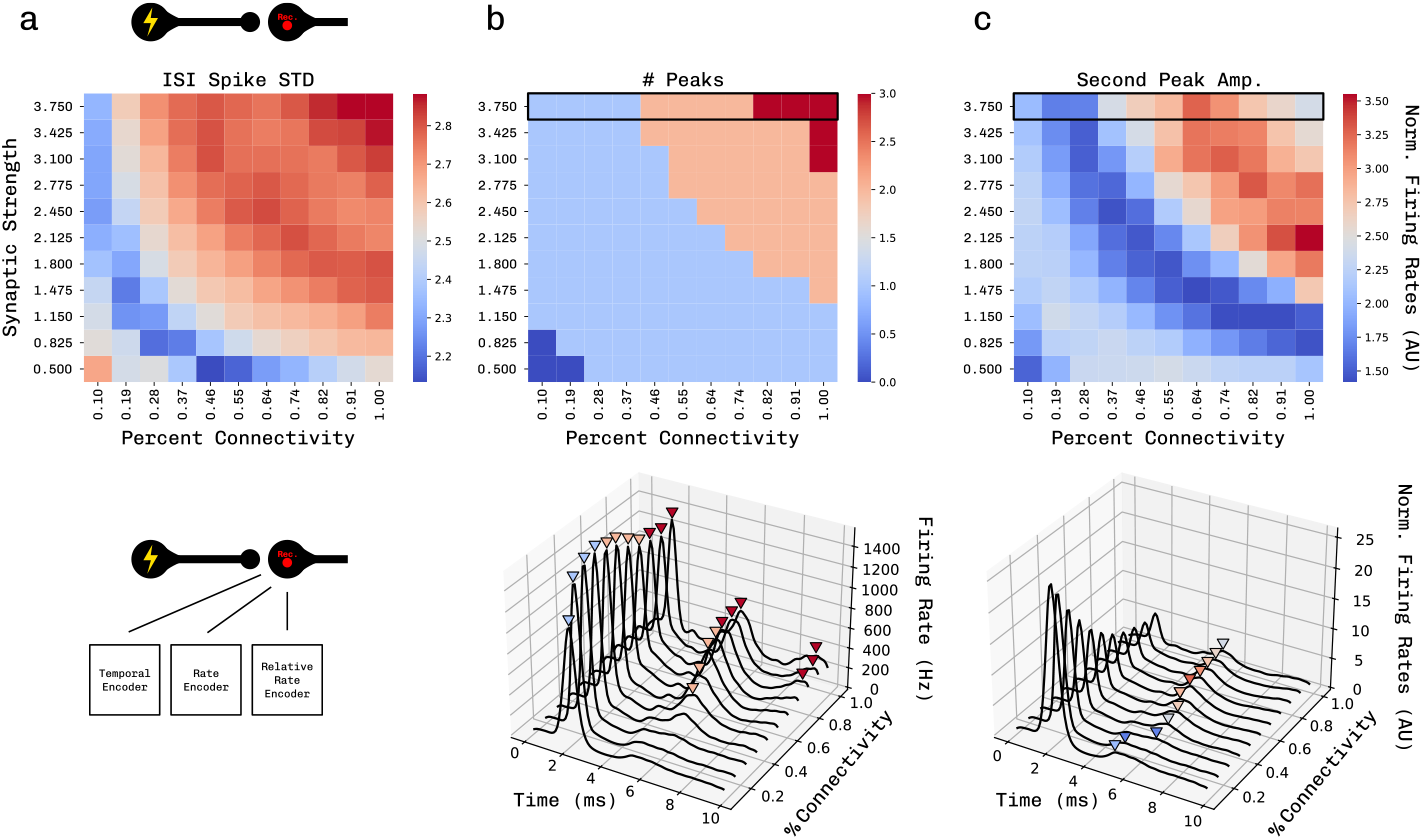
Properties of the inter-stimulus interval in a feed forward network. In all plots, the x-axis is connection probability, and the y-axis is synapse strength. **a)** The feature is spike synchrony (standard deviation). **b)** The features is number of peaks that surpass 250 *Hz*. **c)** Amplitude of the second peak response in the ISI normalized to the baseline, pre-stimulus, firing rate. Below **b) & c)** are example traces of instantaneous ISI firing rates as connectivity is varied, with maximal synapse strength.

**Fig 7.**
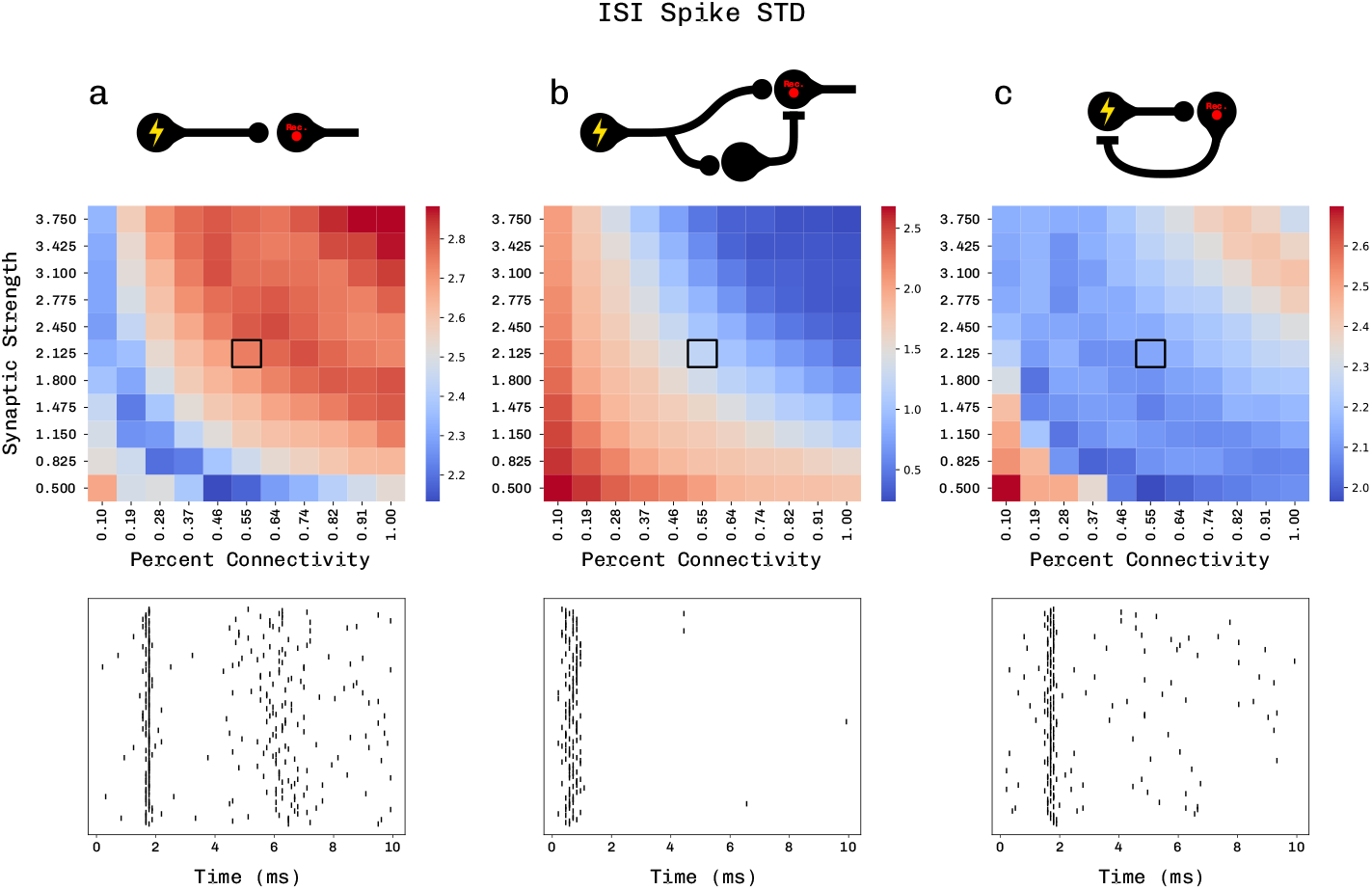
Circuit motif influence. Inter-stimulus spike synchrony with respect to connection probability on the x-axis, and synapse strength on the y-axis. **a)** Synchrony of the ISI in a feed-forward motif, as in Figure 5. **b)** Synchrony of the ISI in a network with lateral inhibition. **c)** Synchrony of the ISI in a network with recurrent inhibition.

**Fig 8.**
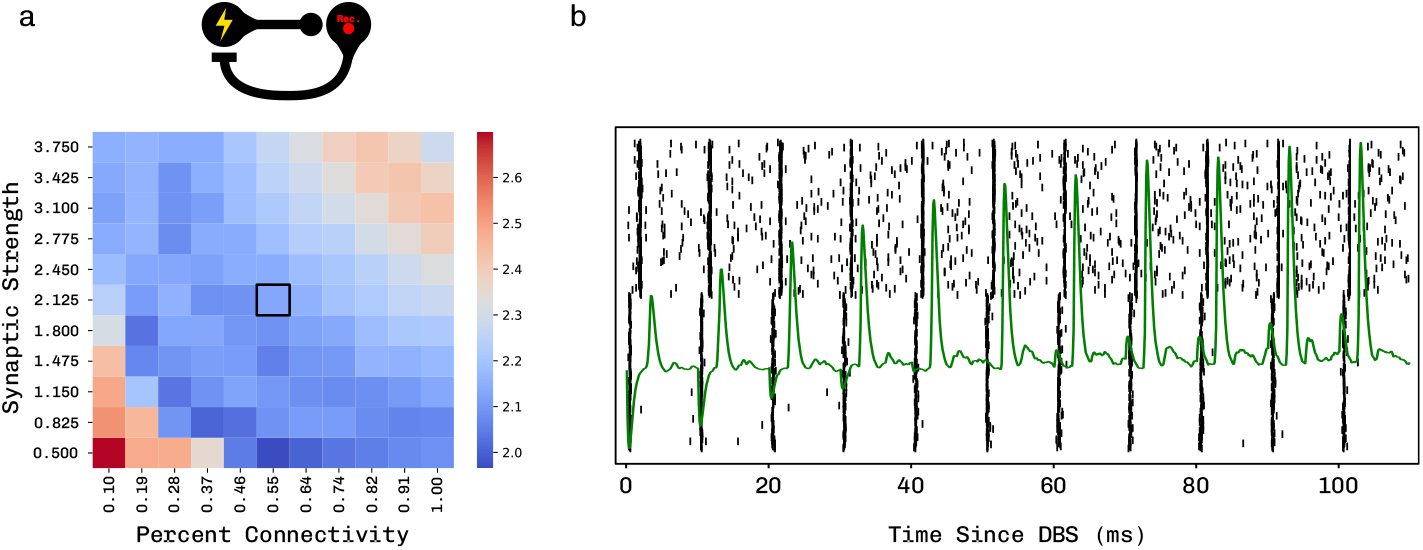
Evoked Field Oscillations. This figure depicts how certain motifs may give rise to oscillations that are a function of downstream connectivity that is modulated by DBS to give rise to an oscillatory or patterned response. a) is the feature space the model was simulated in. b) is the raster (black) and LFP (green) of the first 10 pulses to high frequency DBS

**Fig 9.**
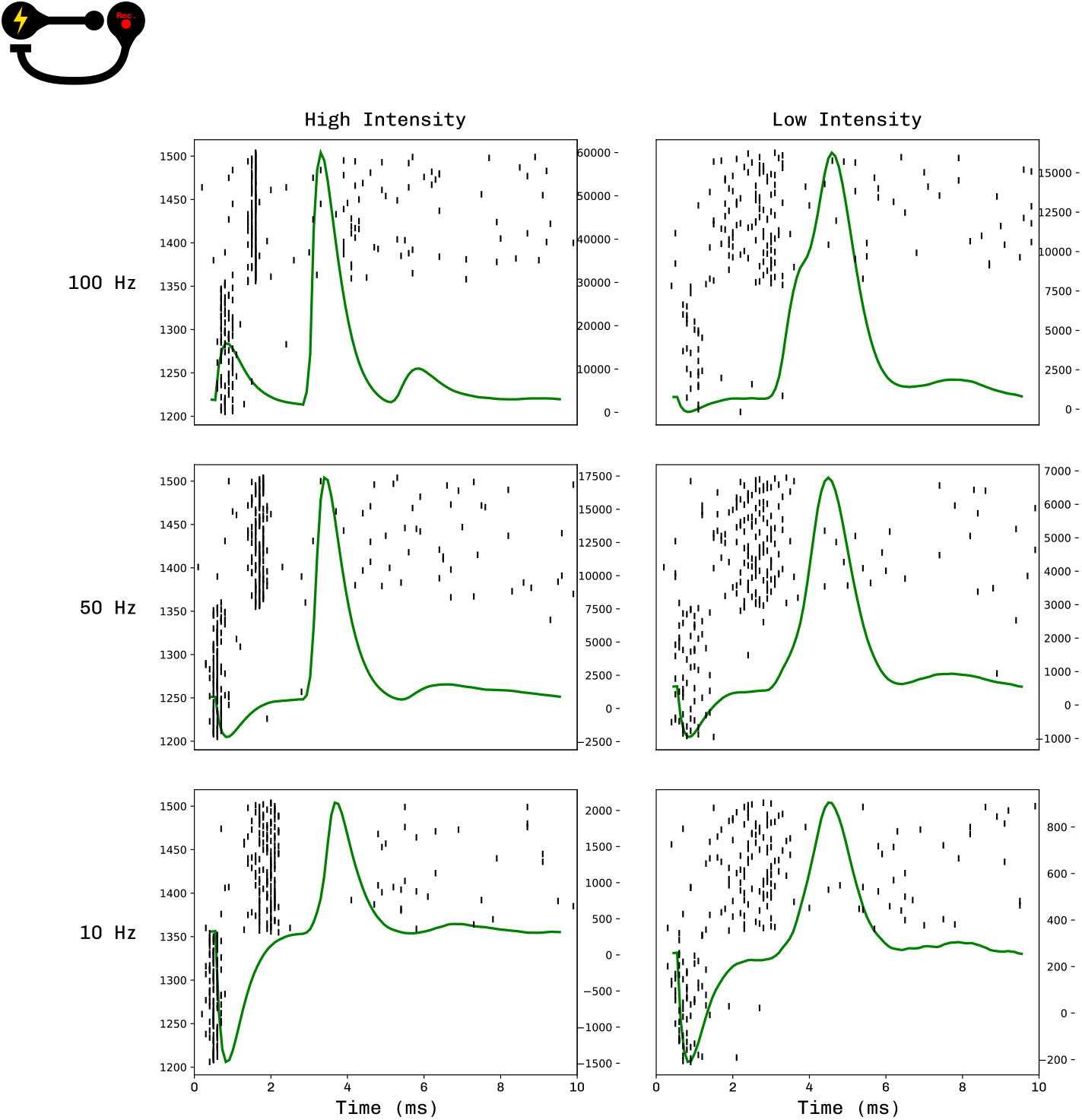
Evoked Field Oscillations Modulated by Stimulation Parameters. How delayed local evoked potentials (DLEPs) are effected by different stimulation parameters such as stimulation intensity (column)1 and frequency (row).

Computational models of deep brain stimulation have been growing in popularity for fibre recruitment, especially with utilization of volume conductor models and similar to estimate the volume of activated tissue (VAT) [35, 36]. These techniques function depend on a detailed characterization of the intersecting fibres in order to estimate what fibres might interact with each DBS pulse. While many trajectories have been characterized in good detail, such as the hyperdirect pathway [11], many tracts such as those between the basal ganglia nuclei are only roughly estimated by probabilistic tractography [37, 38]. These probabilistic estimates of connectivity within the basal ganglia are unable to meaningfully distinguish subthalamic nucleus-globus pallidus externus (STN-GPe) and STN-globus pallidus internus (STN-GPi), which are thought to play an important role, especially the STN-GPe connectivity in the clinical efficacy of treatment of Parkinson’s disease and Dystonia [13, 34]. To allow for investigating the differential fibre recruitment profiles of these precise fibres we set out to develop this framework.

### Limitations

While this method allows for a drop-in implementation of DBS into existing networks, there are many factors that it does not account for without careful consideration. The calculation of the number of recruited efferent, and afferent fibres is dependent on the user to determine, from methods such as the volume of activated tissue. This method also does not account for any direct sub-threshold changes to dynamics that DBS may induce, which may interact with the soma leaving it in a more or less excitable state which this method fails to capture. Further, this method can introduce a great degree of computational complexity in certain simulators, such as NEST and Brian, as all connections become interposed with parrot neurons, requiring then more spike propagation, which can substantially slow simulation time.

## Conclusion

With the growing efforts to optimize and better understand the therapeutic mechanism of DBS, fibre specific recruitment is thought to play an integral role. Herein we present a standard computational framework to interrogate fibre specific activations, in a physiologically constrained computationally tractable manner. The model allows for the integration of known physiological properties, such as those of activation latencies and dose responses, while allowing for exploration of fibre recruitment properties for less defined fibres, e.g. intra-basal-ganglia fibres. The model of stimulation generalizes to any spiking network stimulation by injecting spike times directly at synaptic elements. This direct integration allows for features such as short term synaptic plasticity to be included in the effect of DBS. To showcase the tractability and generalization of this approach we generated circuits of different structural and temporal motifs and showed how differentially DBS affected these networks.

## Acknowledgments

This section is intended only for general acknowledgements and thanks. Any information related to funding, data availability, author contributions, etc. should be entered directly into their dedicated fields in the PLOS Editorial Manager submission system, which will then be incorporated into the appropriate section in your article during the production process.

